# SmCCNet 2.0: A Comprehensive Tool for Multi-omics Network Inference with Shiny Visualization

**DOI:** 10.1101/2023.11.20.567893

**Authors:** Weixuan Liu, Thao Vu, Iain Konigsberg, Katherine Pratte, Yonghua Zhuang, Katerina Kechris

## Abstract

**Summary:** Sparse multiple canonical correlation network analysis (SmCCNet) is a machine learning technique for integrating omics data along with a variable of interest (e.g., phenotype of complex disease), and reconstructing multi-omics networks that are specific to this variable. We present the second-generation SmCCNet (SmCCNet 2.0) that adeptly integrates single or multiple omics data types along with a quantitative or binary phenotype of interest. In addition, this new package offers a streamlined setup process that can be configured manually or automatically, ensuring a flexible and user-friendly experience.

**Availability:** This package is available in both CRAN: https://cran.r-project.org/web/packages/SmCCNet/index.html and Github: https://github.com/KechrisLab/SmCCNet under the MIT license. The network visualization tool is available at https://smccnet.shinyapps.io/smccnetnetwork/.

## 1 Background

Advancements in sequencing and mass spectrometry technologies have allowed access to extensive -omics data sets such as transcriptomics, proteomics, and metabolomics, which allows the integration of different omics data to gain biological insights into complex diseases [1]. Multi-omics network inference is a technique for integrating mul-tiple -omics data sets to infer molecular interactions with respect to trait(s) of complex disease and gain insights into associated biological processes [2]. Recent multi-omics network inference methods include knowledge-guided multi-omics network inference (KiMONo) [3] and Biomarker discovery using Latent Variable Approaches for Omics Studies (DIABLO) [4].

Sparse multiple Canonical Correlation Network Analysis (SmCCNet) is a canonical correlation-based integration method that reconstructs phenotype-specific multi-omics networks [5]. It has been applied to different multi-omics integration tasks such as extracting protein-metabolite networks [6] and mRNA-miRNA networks [7] associated with disease phenotypes. The algorithm is based on a sparse multiple canonical analysis (SmCCA) [8] for *T* omics data *X*_1_, *X*_2_, …*X*_*T*_ and a quantitative phenotype *Y* measured in the same subjects. SmCCA finds the canonical weights *w*_1_, *w*_2_, …, *w*_*T*_ that maximize the (weighted or unweighted) sum of pairwise canonical correlations between *X*_1_, *X*_2_, …, *X*_*T*_ and *Y*, under sparsity constraints in Equ 1. In SmCCNet, the sparsity constraint functions *P*_*t*_(*·*), *t* = 1, 2, …, *T*, are the least absolute shrinkage and selection operators (LASSO) [9]. The weighted version corresponds to *a*_*i,j*_, *b*_*i*_ (also called scaling factors), which are not all equal; the unweighted version corresponds to *a*_*i,j*_ = *b*_*i*_ = 1 for all *i, j* = 1, 2, …, *T*, where *a*_*i,j*_ are for between -omics relationships, while *b*_*i*_ is for the single omics and phenotype relationship.

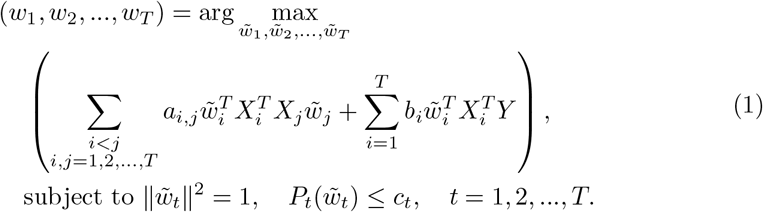

The sparsity penalties *c*_*t*_ influence how many features will be included in each subnetwork. With pre-selected sparsity penalties, the SmCCNet algorithm creates a network similarity matrix based on SmCCA canonical weights from repeatedly sub-sampled omics data and the phenotype and then finds multi-omics modules that are relevant to the phenotype. The subsampling scheme improves network robustness by analyzing subsets of omics features multiple times and forms a final similarity matrix by aggregating results from each subsampling step.

The first version of SmCCNet could only analyze two omics data types with a quantitative phenotype, did not scale well for large data sets, used simple network summarization that did not incorporate d network structure, required a complicated manual setup and limited visualization. To enhance the practical utility of SmCCNet with improved or novel methods and functionalities, we have rewritten and upgraded the software to flexibly accommodate one or more omics data types, as well as a binary phenotype. We also created an automated end-to-end pipeline that obtains the final network result with just a single line of code. Other improvements include a data pre-processing pipeline, improved computational efficiency, an online RShiny application for network visualization, and the storage of accessible and reproducible network analysis results in a user-specified directory. The full list of changes for SmCCNet 2.0 is listed below:

- Multi-omics SmCCNet with quantitative phenotype allows integration of more than two -omics data **(improved functionality, section 2.1.1)**.
- Novel hybrid SmCCNet algorithm with binary phenotype **(novel functionality and algorithm, section 2.1.2)**.
- Single-omics implementation of SmCCNet algorithm with either quantitative or binary phenotype **(novel functionality and algorithm, section 2.1.3)**.
- Novel model-wise optimal scaling factors selection algorithm for multi-omics SmCCNet **(novel functionality and algorithm, section 2.2)**.
- Implementation of NetSHy network summarization method to summarize network based on network topology **(novel functionality, section 2.3)**.
- Subnetwork pruning algorithm to reduce the size of multi-omics network **(novel functionality and algorithm, section 2.3)**.
- New subnetwork visualization RShiny application for multi-omics interaction network visualization **(novel functionality, section 2.4)**.
- Fast Automated SmCCNet conducts end-to-end pipeline with a single line of code and further improves the algorithm speed **(novel functionality, section 2.5)**.
- New -omics data preprocessing pipeline to filter out features with low variability, regress out clinical covariates, and center/scale **(novel functionality, described in section 2)**.
- Simpler coding setup to run SmCCNet manually and boost the algorithm speed by 100-1000x **(improved functionality, described in section 2)**.

## 2 Implementation

Before running SmCCNet, the user can apply a streamlined function to preprocess the omics data, including filtering features with low Coefficient of Variation (CoV), centering and scaling each molecular feature, and regressing out effects from covariates. The data preprocessing pipeline can be implemented by using the *dataPreprocess()* function. The end-to-end pipeline of SmCCNet takes in any number of molecular profiles (omics data) and either a quantitative or binary phenotype, and outputs the single/multi-omics subnetwork modules that are associated with the phenotype. The general workflow of SmCCNet is shown in Figure 1.

**Fig. 1.**
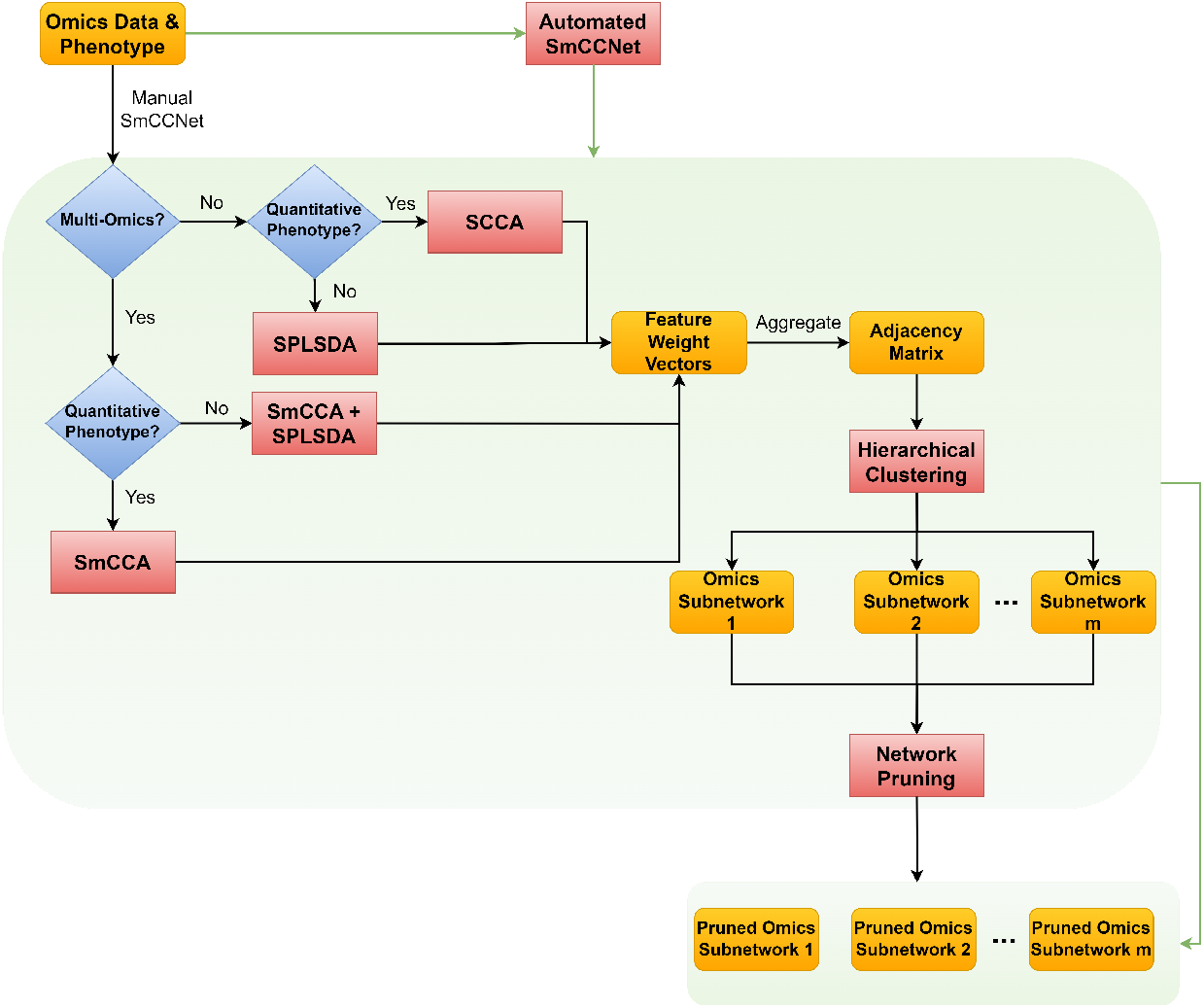
SmCCNet workflow. The workflow of the second-generation SmCCNet, which takes in single-/multi-omics data with quantitative/binary phenotype and outputs multiple pruned omics subnetwork modules. Steps in the box represent the stepwise process of executing the SmCCNet, and if automated SmCCNet is used, all these steps within the box will be executed automatically.

### 2.1 Number of Omics Data and Phenotype Modality

In general, SmCCNet consists of the following steps (See Figure 2, 3, and 4):

**Fig. 2.**
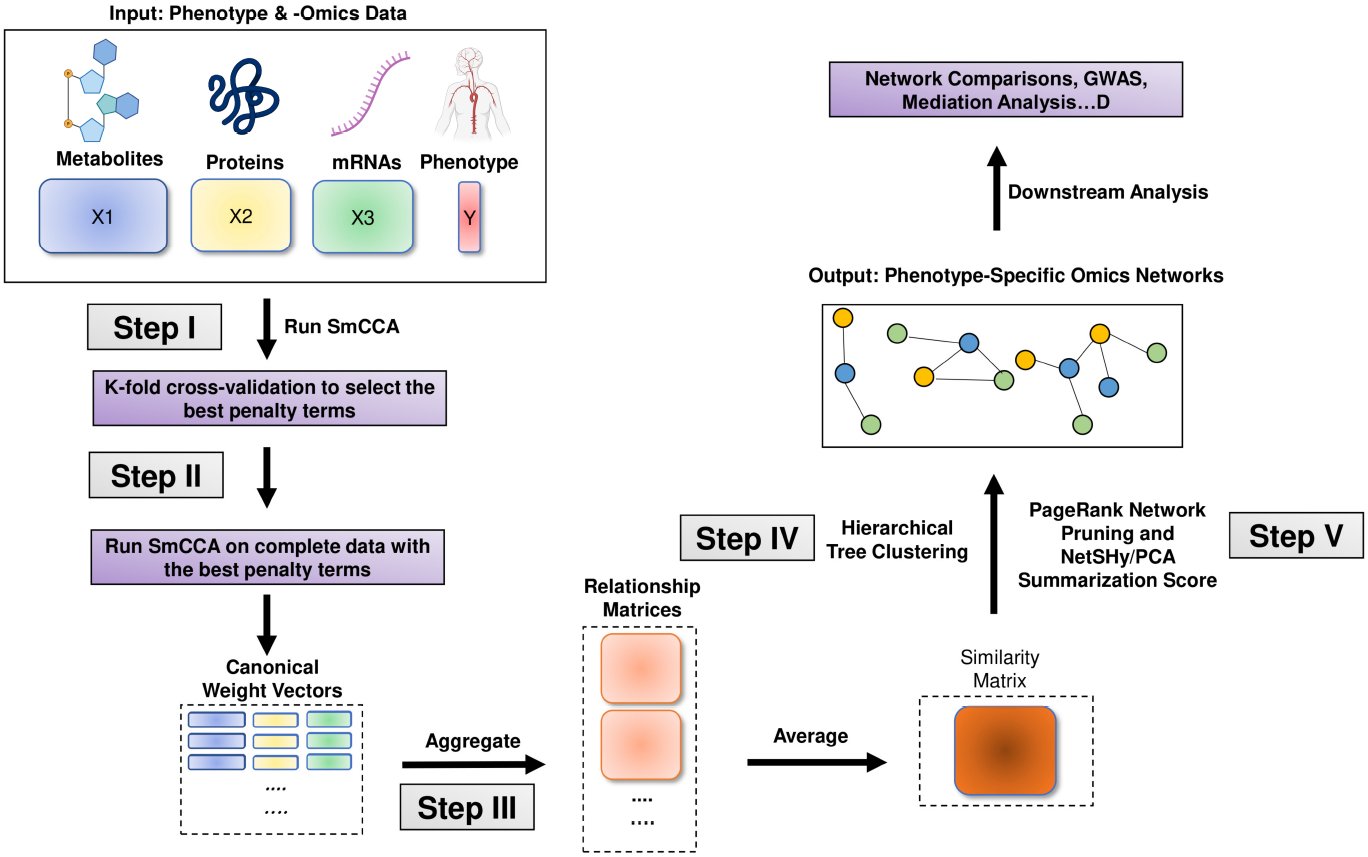
Multi-Omics SmCCNet with Quantitative Phenotype workflow. T he w orkflow of the second-generation SmCCNet with multi-omics data and quantitative phenotype.

**Fig. 3.**
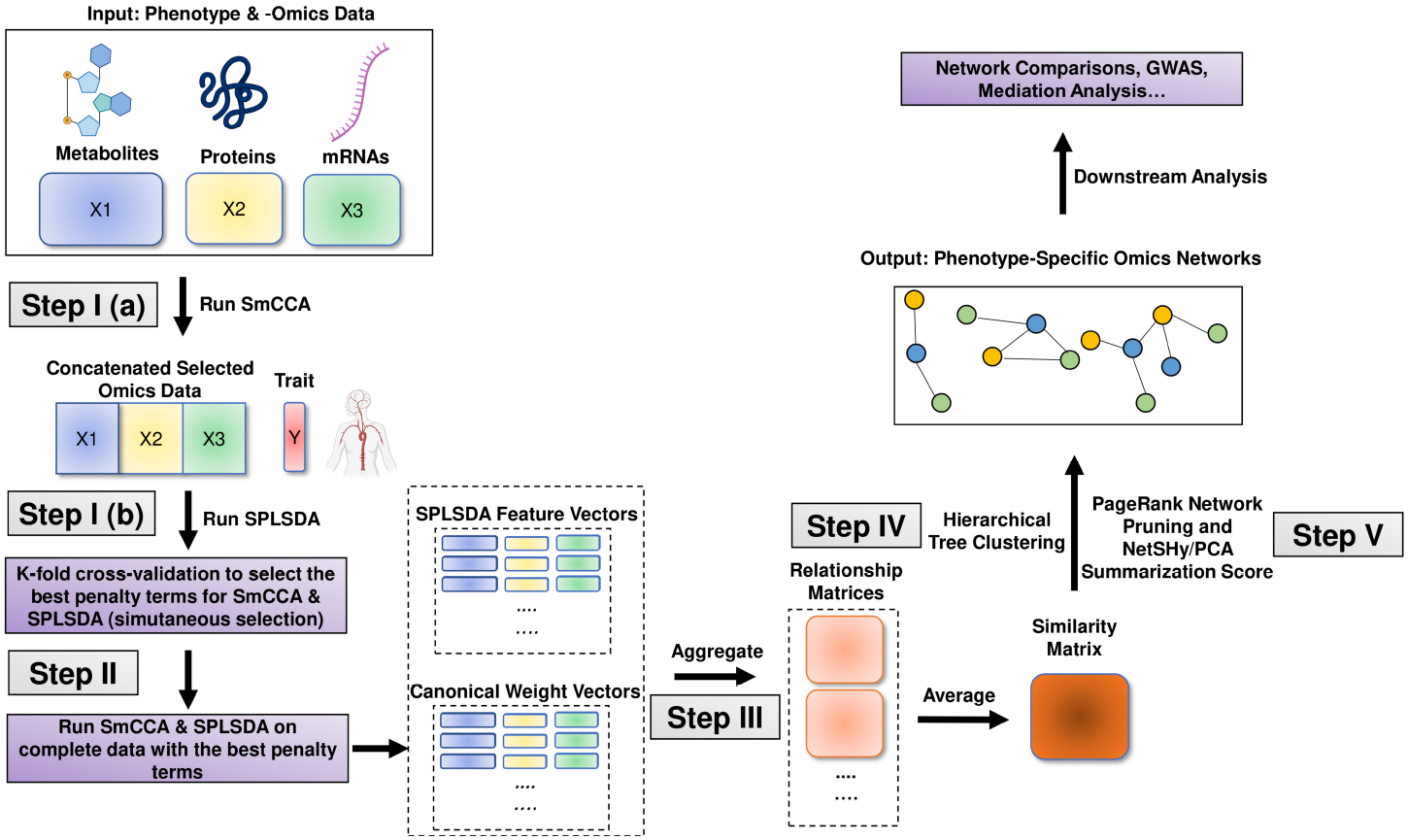
Multi-Omics SmCCNet with Binary Phenotype workflow. The workflow of the second-generation SmCCNet with multi-omics data and binary phenotype.

**Fig. 4.**
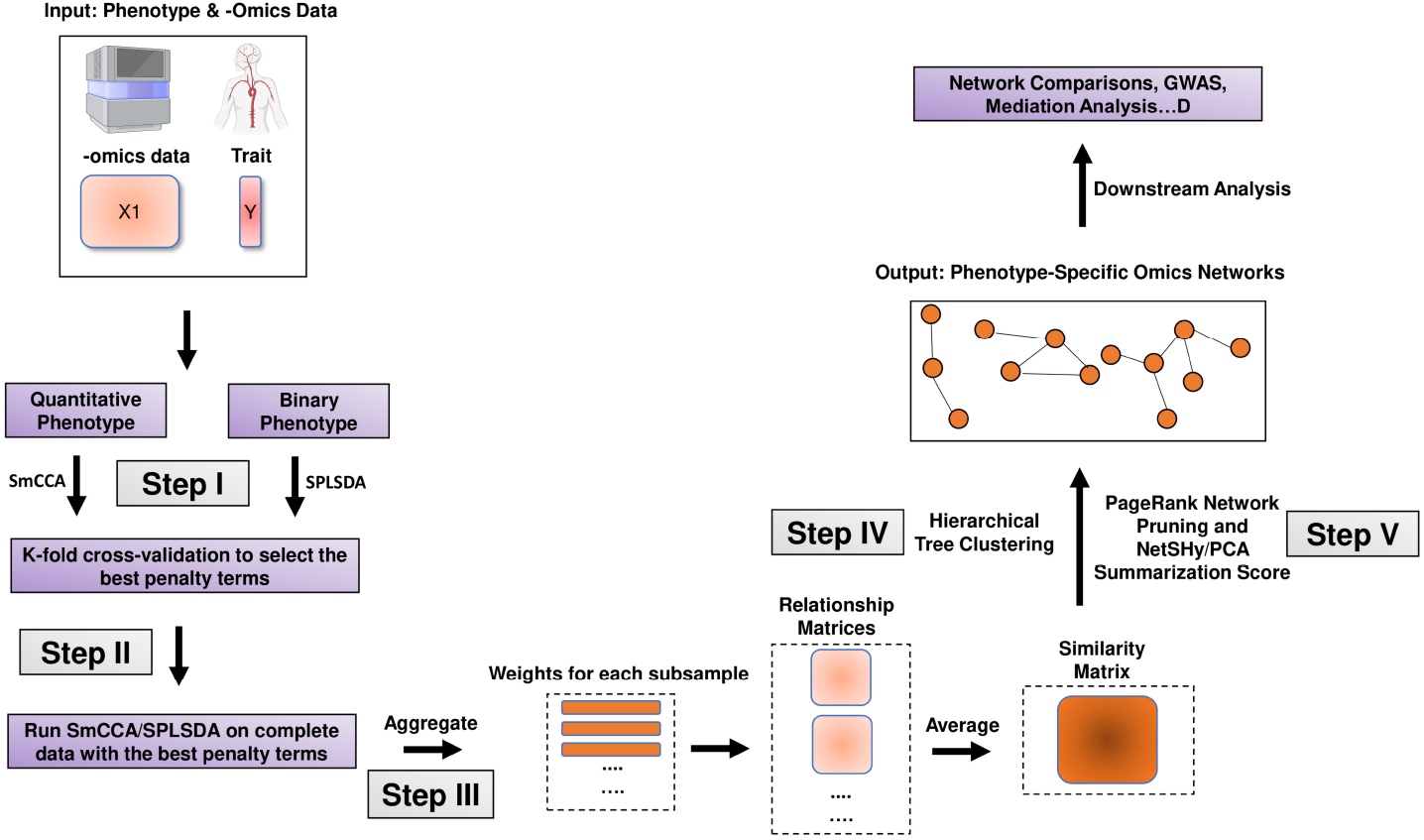
Single-Omics SmCCNet with Quantitative/Binary Phenotype workflow. The work-flow of the second-generation SmCCNet with single-omics data and quantitative/binary phenotype.

- Step I: Determine SmCCA/SPLSDA sparsity penalty parameters. The user can select the penalties for feature selection based on prior knowledge. Alternatively, the user can pick sparsity penalties based on a K-fold cross-validation (CV) procedure that minimizes the total prediction error (Figure 1). The K-fold CV procedure enhances the robustness of selected penalties when generalizing to similar independent omics data sets.
- Step II: Randomly subsample omics features, apply SmCCA/SPLSDA with chosen penalties and compute a canonical weight vector for each subsample. Repeat the process many times.
- Step III: Compute feature similarity matrix based on canonical weight matrix.
- Step IV: Apply hierarchical tree cutting to the similarity matrix to simultaneously identify multiple subnetworks.
- Step V: Apply network pruning algorithm to prune each subnetwork obtained from Step IV and Visualize the -omics network with an RShiny application (https://smccnet.shinyapps.io/smccnetnetwork/) or Cytoscape.

Steps III to V remain consistent across all scenarios, regardless of the number of omics data types used and the phenotype modality involved. However, Steps I and II differ depending on the specific scenario. Below, we provide detailed descriptions for Steps I and II for each scenario:

#### 2.1.1 Multi-omics SmCCNet with Quantitative Phenotype

If multi-omics data is used with quantitative phenotype, same as SmCCNet 1.0, we implement the SmCCA algorithm (Equation 1) for feature selection and network construction, which is achieved by using *getRobustWeightsMulti()*. In step I, we use *k*-fold cross-validation to determine the optimal penalty parameters based on the loss function. In SmCCNet 1.0, the loss function is defined to be the prediction error, which is defined as follows:

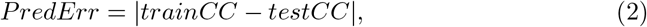

where *trainCC* and *testCC* are defined a s t he t raining c anonical c orrelation and testing canonical correlation respectively. In SmCCNet 2.0, the loss function is defined to be the scaled prediction error:

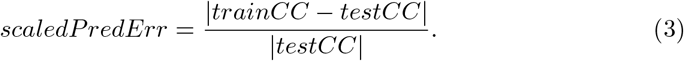

Compared to prediction error, the scaled prediction error aims not only to minimize the discrepancy between the training and testing canonical correlations but also to maximize the testing canonical correlation. This approach effectively prevents the selection of penalty parameters that could result in extremely low testing canonical correlations.

In Step II, to enhance the robustness of the multi-omics network, we employ a subsampling algorithm ^1^. This algorithm selects only a fraction of the molecular features from each molecular profile during each iteration of subsampling. The complete workflow of multi-omics SmCCNet with quantitative phenotype is shown in Figure 2. The detailed implementation can be found in the package vignette ^2^.

#### 2.1.2 Multi-omics SmCCNet with Binary Phenotype

We developed the hybrid algorithm between Sparse Partial Least Squared Discriminant Analysis (SPLSDA) [10] and SmCCA for feature selection and network construction, which is achieved by using *getRobustWeightsMultiBinary()*. SPLSDA is a two-step approach to implement the partial least squared algorithm with the binary outcome, which is a combination of partial least squared and logistic regression.

First, SmCCA (Sparse Multiple Canonical Correlation Analysis) is applied without involving phenotype data to filter molecular features and identify those that are interconnected. Next, SPLSDA (Sparse Partial Least Squares Discriminant Analysis) is employed on the chosen features across all molecular profiles to determine which features are associated with the phenotype. Finally, the canonical weights obtained from SmCCA and the feature importance weights from SPLSDA are combined into a weighted average, providing a consolidated measure of each feature’s relevance.

To select the optimal penalty parameters, *k*-fold cross-validation is implemented on the complete hybrid algorithm to evaluate penalty terms on SmCCA and SPLSDA simultaneously. In SmCCNet 2.0, various metrics can be used to evaluate each set of penalty parameters, which include prediction accuracy, AUC score, precision, recall, and F1 score.

After the optimal penalty terms are selected, the hybrid method is run on the complete dataset, and same as regular SmCCNet, we use the subsampling scheme to ensure the robustness of the multi-omics network.

The complete workflow of multi-omics SmCCNet with quantitative phenotype is shown in Figure 3. Consider *X*_1_, *X*_2_, …, *X*_*T*_ as *T* omics datasets, and *Y* as the phenotype data. The hybrid SmCCNet algorithm with binary phenotype is defined as follows:

- **Stage 1: Run Weighted/Unweighted Sparse Multiple Canonical Correlation Analysis (SmCCA):** This is performed on *X*_1_, *X*_2_, …, *X*_*T*_ (excluding phenotype data). The output is canonical weight vectors (with nonzero entries, zero entries are filtered) 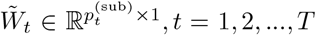, which represent the omics-omics connections. In this step, we filter out features that have no connections with other features, which helps reduce dimensionality. Note that we tend to set relaxed penalty terms for this step to include as many omics features as possible to increase the performance of the classifier in the next step.
- **Stage 2: Subset Omics Data:** Each dataset *X*_1_, *X*_2_, …, *X*_*T*_ is subsetted to include only omics features selected in Step 1, calling subsetted data 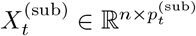
- **Stage 3: Concatenate and Run Sparse Partial Least Squared Discriminant Analysis (SPLSDA):** The subsetted datasets 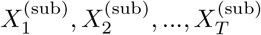 are concatenated into 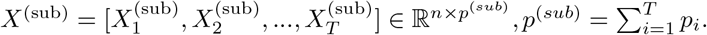 The SPLSDA algorithm is then run to extract *R* latent factors and a projection matrix, by default, *R* is set to 3. The projection matrix is defined as 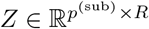. Latent factors are defined as *L* = [*r*_1_, *r*_2_, …, *r*_*R*_] = *X*^(sub)^ *· Z ∈* R^*n×R*^.
- **Stage 4: Aggregate Latent Factors:** The *R* latent factors are aggregated into one using logistic regression, defined by logit(*Y*) = *α*_1_*r*_1_ + *α*_2_*r*_2_ + … + *α*_*R*_*r*_*R*_. Feature weights are given by aggregation of the projection matrix from Sparse PLSDA 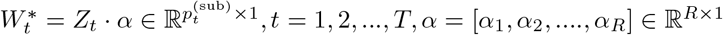, where *Z*_*t*_ is the subset of projection matrix *Z* such that it only includes features from the *t*th omics data.
- **Stage 5: Normalize and Calculate Final Canonical Weight:** The feature

weights 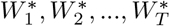 based on SPLSDA are normalized to have an L2 norm of 1. Let *γ*_1_ and *γ*_2_ be two scalars representing the strength of omics-omics and omics-phenotype connections, respectively. The final canonical weight is obtained by combining the canonical weight from step 1 and the feature weight from the classifier from step 4: 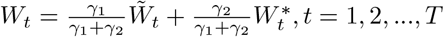

#### 2.1.3 Single-omics SmCCNet with Quantitative/Binary Phenotype

In multi-omics SmCCNet, the between-omics interaction is taken into account. However, in the single-omics setting, this is no longer considered. Therefore, we developed two functions separately to tackle single-omics analysis (Figure 4). If a quantitative phenotype is used, then sparse canonical correlation analysis (SCCA) is used to construct the global network by using the function *getRobustWeightsSingle()*; if a binary phenotype is used, then both stage 3 and 4 of the hybrid algorithm above are used with SPLSDA algorithm by using the function *getRobustWeightsSingleBinary()*. For more information about the single-omics SmCCNet pipeline setup, runnable examples are provided in the package vignette. In addition, this pipeline has been applied to the proteomics network analysis to identify the single-omics networks associated with pulmonary function and smoking behavior [11].

### 2.2 Scaling Factor Determination

The scaling factors (*a*_*i,j*_ and *b*_*j*_) in Equation 1 can be supplied to prioritize the correlation structure of interest in Steps I and II of the SmCCNet Pipeline. Users can choose to supply their own choice of scaling factors or select them with the model-based approaches. We provide three different methods for selecting the scaling factors.

#### 2.2.1 Prompt to Define Scaling Factors

If a user is able to supply the scaling factors for the model based on prior knowledge, an interactive function *scalingFactorInput()* can be used to enter scaling factors manually for each pairwise correlation. For instance, when entering *scalingFactorInput(c(‘mRNA’,’miRNA’, ‘phenotype’))*, three sequential prompts will appear, requesting the scaling factors for mRNA-miRNA, mRNA-phenotype, and miRNA-phenotype relationships, respectively.

#### 2.2.2 Pairwise Correlation to Select Scaling Factors with Automated SmCCNet

As an alternative, the pairwise correlation between each pair of omics data can be used to set the scaling factors. For this option, SCCA is run with a stringent penalty pair. The resulting canonical correlation will be treated as the between-omics scaling factor, while a scaling factor of 1 will be used for the omics-phenotype relationship. In addition, we introduce another parameter called the shrinkage factor to prioritize either the omics-omics relationship or the omics-phenotype relationship. For example, in a multi-omics analysis with two omics data, if the omics-omics correlation is 0.8 by SCCA, and the shrinkage parameter is 2, then the final scaling factors are set to (*a, b*_1_, *b*_2_) = *c*(0.4, 1, 1), where *a* are the between-omics relationship and *b*’s are the omics-phenotype relationships. This method is currently implemented in the automated SmCCNet approach.

#### 2.2.3 Cross-Validation to Select Scaling Factors

The approach employs cross-validation to identify optimal scaling factors, illustrated using two omics types as an example. Initially, candidate sets of scaling factors are generated with all omics-omics scaling factors set to 1, and omics-phenotype scaling factors adjusted so their sum equals 1 for comparability. For instance, scaling factors (*a*_1,2_, *b*_1_, *b*_2_) must fulfill the condition *a*_1,2_ + *b*_1_ + *b*_2_ = 1. A nested grid search strategy is then applied to simultaneously optimize the scaling factors and penalty parameters. Within this framework, as different sets of scaling factors are evaluated, the optimal penalty parameters are selected. For each candidate set of scaling factors, the optimal sparse penalty parameters (denoted as *l*1, *l*2) are identified via *k*-fold cross-validation. The evaluation metric’s value corresponding to these parameters is recorded, which is associated with the optimal penalty parameters for each candidate set. This process is repeated across all potential combinations of scaling factors. The set of scaling factors yielding the best performance, according to the chosen evaluation metric, is selected as the optimal set, together with its associated optimal penalty parameters. Given the exponential increase in possible scaling factor combinations with more than three types of -omics data, the use of the automated SmCCNet algorithm is recommended for selecting optimal scaling factors in analyses involving larger numbers of -omics data types.

### 2.3 Network Clustering and Pruning

The adjacency matrix is formed by taking the outer product of the canonical weights. After obtaining the adjacency matrix, hierarchical clustering [12] is implemented to partition molecular features into different network modules, and this is achieved by using the function *getAbar()*.

The objective of Step V is to prune the network by removing features (nodes) that have no/little contribution to the subnetwork using a network summarization score of Principal Component Analysis (PCA) [13] or network summarization via a hybrid approach leveraging topological properties (NetSHy) [14] to produce a densely connected pruned subnetwork that maintains a high summarization correlation with respect to the phenotype (Figure 5). Initially, the network features are ranked based on their PageRank scores [15]. Beginning with a user-defined minimum baseline network size, the method iteratively includes additional features, evaluating the summarization correlation with respect to both the phenotype and the baseline network at each step until reaching the optimal subnetwork size. The network pruning step is achieved by implementing the function *networkPruning()*, and the step-by-step description is given as follows:

**Fig. 5.**
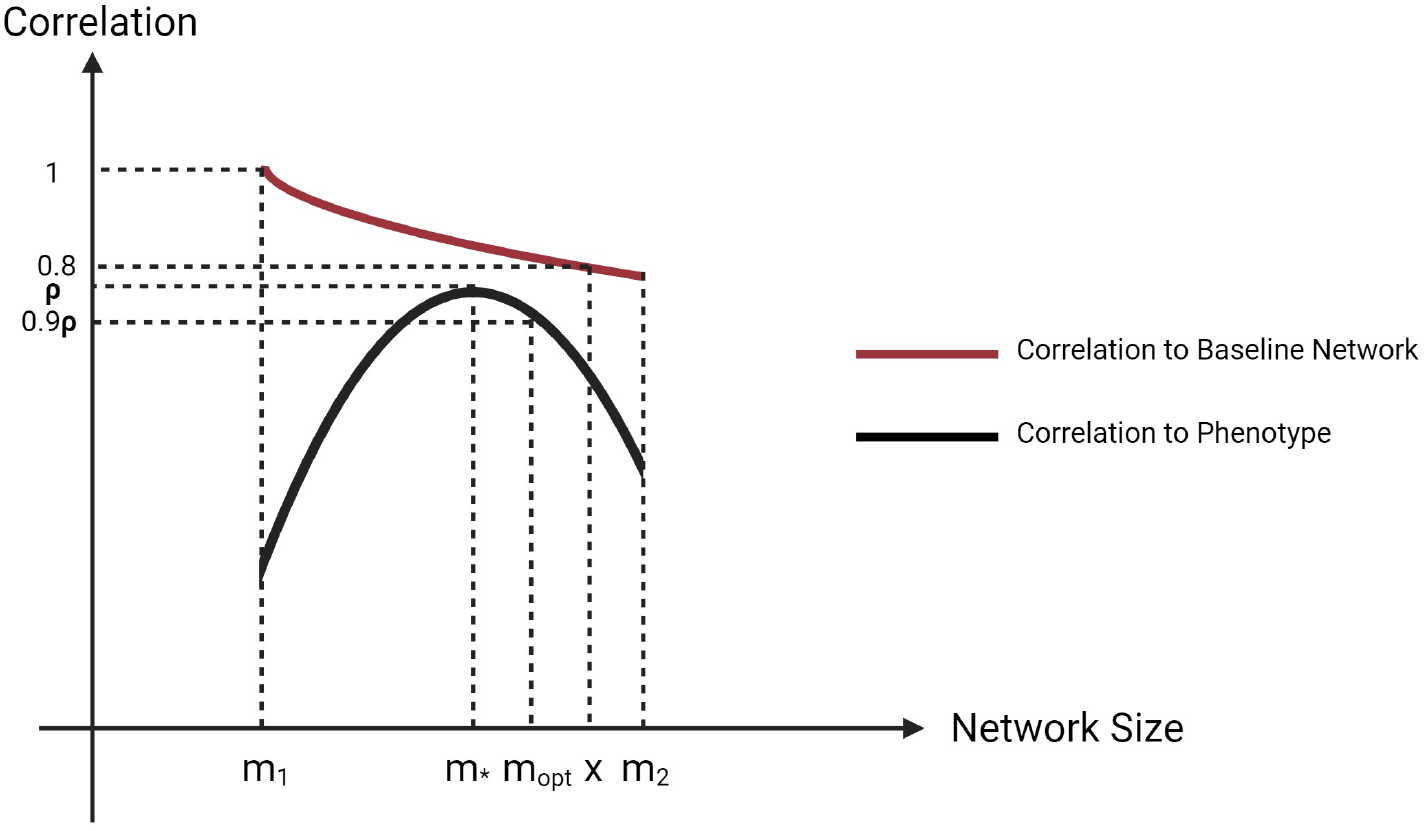
Conceptual figure of network pruning algorithm with the y-axis to be NetSHy/PCA summarization score’s correlation to phenotype (black) or baseline network at *m*_1_ (red). *m*_*∗*_ is the network size with the highest correlation to phenotype. *x* (*x* ¿ *m*_*∗*_) is the maximum network size that has a least 0.8 correlation to the baseline network at *m*_1_. *m*_*opt*_ corresponds to the optimal network size.

- Calculate PageRank score for all molecular features in the unpruned network and rank them according to PageRank score.
- Start from minimally possible network size *m*_1_, iterate the following steps until reaching the maximally possible network size *m*_2_ (defined by users):
  – Add one more molecular feature into the network based on node ranking, then calculate NetSHy/PCA summarization score (PC1, PC2, PC3) for this updated network.
  – Calculate the correlation between this network summarization score and phenotype for the current network size *i ∈* [*m*_1_, *m*_2_], and only use the PC with the highest correlation (determined by absolute value) w.r.t. phenotype, define this correlation as *ρ*_(*i,pheno*)_.
- Identify network size *m*_*∗*_ (*m*_*∗*_ *∈* [*m*_1_, *m*_2_]) with *ρ*_(*m∗,pheno*)_ being the maximally possible summarization score correlation w.r.t. phenotype (determined by absolute value).
- Treat *m*_*∗*_ as the new baseline network size, let *ρ*_(*m∗,i*)_ be the correlation of summarization score between network with size *m*_*∗*_ and network with size *i*. Define *x* to be the network size (*x ∈* [*m*_*∗*_, *m*_2_]), such that *x* = max {i|(*i ∈* [*m*_*∗*_, *m*_2_])&(|*ρ m*_*∗*_,*i*) | *>* 0.8)}.
- Between network size of *m* and *x*, the optimal network size *m*_*opt*_ is defined to be the maximum network size such that |*ρ m*_(*opt,pheno*)_ | *≥* 0.9 *·* |*ρ*_(*m,pheno*)_|.

### 2.4 Network Visualization

The SmCCNet pipeline saves the final subnetwork information in a .Rdata file, which does not include data for network visualization. To enable the translation of this .Rdata file into a visual representation of the network, we have developed an R Shiny application, accessible at https://smccnet.shinyapps.io/smccnetnetwork/ (Figure 6). This application provides a user-friendly platform for visualizing single or multi-omics networks, utilizing subnetworks created and stored by SmCCNet. To begin, users can easily generate an example network visualization by simply clicking the ‘Plot Network’ button, with no need for uploading any data. For more customized visualizations, users can upload a .Rdata file with the naming convention ‘size a net b.Rdata’, where ‘a’ represents the pruned network size, and ‘b’ indicates the network module index following hierarchical clustering.

**Fig. 6.**
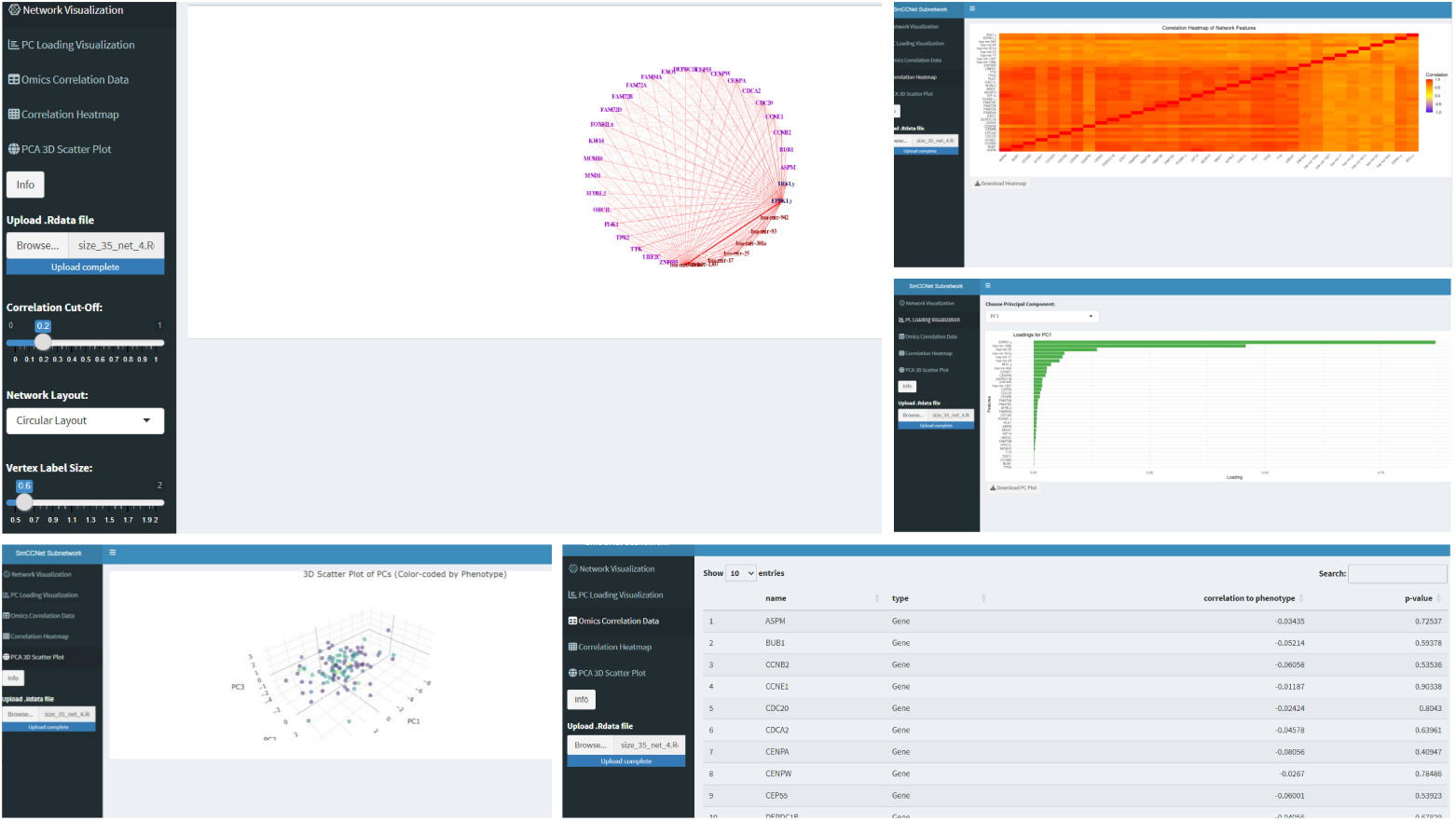
Example R Shiny Interface for Network Information Visualization. The example interface for network visualization. The users can upload network .*Rdata* file to the application, and tune the visualization parameters to obtain the optimal visual representation of the -omics network. Other network-relevant information (PC loading bar graph, correlation heatmap, 3-D subject plot, feature-phenotype correlation table) are also demonstrated

To refine the network visualization, the application offers several adjustable parameters. The Correlation Cut-Off slider allows users to filter network edges based on correlation values, enabling a focus on more significant connections. The Network Layout drop-down menu presents different layout options, which facilitates the selection of the preferred visual arrangement for the network. Users can also adjust the sizes of vertex labels and vertices through the respective Vertex Label Size and Vertex Size sliders. Moreover, the Edge Intensity slider provides control over the color intensity and width of the edges. After adjusting these parameters to their satisfaction, users can generate the network visualization by clicking the ‘Plot Network’ button. The ‘Download Plot’ button enables the download of the network visualization as a PDF. Additionally, this application also enables the demonstration of (1) the correlation matrix heatmap between network features; (2) the visualization of PC loadings for the first 3 PCs; (3) The 3-D graph visualizing the distribution of subjects with respect to the first 3 PCs; and (4) the feature-phenotype correlation table can also be shown in the application (see Figure 6).

The application is optimally designed for visualizing final subnetworks of a relatively small size (e.g., *<* 100 nodes). For larger networks, manual adjustments, such as moving nodes to prevent label overlap, are often necessary. In these instances, we recommend users employ Cytoscape [16] for network visualization. Communication between R and Cytoscape is facilitated by the `RCy3`x package [17].

### 2.5 Automated SmCCNet

In this version of the SmCCNet package, we introduce a pipeline known as Automated SmCCNet, which can be implemented with *fastAutoSmCCNet()*. This method streamlines the SmCCNet code and significantly reduces computation time. Users are simply required to input a list of omics data and a phenotype variable. The program then automatically determines whether it is dealing with a single-omics or multi-omics problem, and whether to use CCA or PLS for quantitative or binary phenotypes respectively. For details of how each method is established and how parameters and coefficients are set, we recommend the user to refer to the multi-omics and single-omics vignettes.

Specifically, for multi-omics SmCCNet, if CCA is employed, the program can automatically select the scaling factors (importance of the pair-wise omics or omics-phenotype correlations to the objective function). This is achieved by calculating the pairwise canonical correlation between each pair of omics under the most stringent penalty parameters. The scaling factor for the omics data A, B pair in SmCCA is set to the absolute value of the pairwise canonical correlation between omics A and B divided by the between-omics correlation shrinkage parameter. By default, all scaling factors linked to the phenotype-specific correlation structure are set to 1. In Automated SmCCNet, users only need to provide a BetweenShrinkage parameter, a positive real number that helps reduce the significance of the omics-omics correlation component. The larger this number, the more the between-omics correlation is shrunk.

Moreover, for multi-omics SmCCNet with a binary phenotype, the scaling factor is not implemented. However, the user needs to provide values for *γ*_1_ (omics-omics connection importance) and *γ*_2_ (omics-phenotype connection importance, see multiomics vignette section 5 for detail). The automated SmCCNet program offers a method to calculate *γ*_1_ while setting the value of *γ*_2_ to 1. This is generally done by averaging all the pairwise omics-omics canonical correlations in the multi-omics dataset.

The program can also automatically select the percentage of features subsampled. If the number of features from an omics data is less than 300, then the percentage of feature subsampled is set to 0.9, otherwise, it’s set to 0.7. The candidate penalty terms range from 0.1 to 0.5 with a step size of 0.1 for single/multi-omics SmCCA, and from 0.5 to 0.9 with a step size of 0.1 for single/multi-omics SPLSDA ^3^ (for both omics-omics SmCCA step and omics-phenotype classifier, see section 5 in the multi-omics vignette for details).

The automated version of SmCCNet typically offers a computational speed advantage over the standard manual SmCCNet, primarily due to the heuristic selection of scaling factors and the parallelization of the cross-validation step. This parallelization is achieved through the use of a parallelized map function in *furrr* package [18], significantly improving the computational speed.

## 3 Example

We demonstrate the second-generation SmCCNet utilizing multi-omics data sourced from the Cancer Genome Atlas Program’s (TCGA) project [19] on breast invasive carcinoma (Firehose Legacy). The dataset contains RNA sequencing data with normalized counts, microRNA (miRNA) expression data, and log-ratio normalized reverse phase protein arrays (RPPA) protein data, all procured from tumor samples. After data pre-processing, there are 107 subjects in our final data with 5039 genes, 823 miRNAs, and 175 RPPAs. Furthermore, we regress out age, race, and radiation therapy status from each molecular feature. We provided 2 different examples of using fast automated SmCCNet for multi-omics analysis ^4^. We use survival time as the quantitative phenotype, and survival status as the binary phenotype.

### 3.1 Multi-omics with Quantitative Phenotype

In the TCGA breast cancer example with a quantitative phenotype (survival time), the analysis can be run with the following code (assuming all X (-omics data list) and Y (survival time) are preprocessed):

~~~
result ← fastAutoSmCCNet (X = X, Y = Y,
                          K = 5, subSampNum = 100,
                          DataType = **c** ( ‘Gene’, ‘miRNA’, ‘RPPA’ ),
                          CutHeight = 0.995,
                          summarization = ‘NetSHy’,
                          BetweenShrinkage = 5,
                          seed = 123456)
~~~

In the first phase of the SmCCNet algorithm, 5-fold cross-validation is used to optimize the sparsity penalty for each molecular profile and determine the best scaling factors. We consider the SmCCA penalty parameter for each molecular profile ranging from 0.1 to 0.5, with increments of 0.1, resulting in a total of 125 combinations. The preliminary CCA canonical correlation is 0.960 (gene-miRNA), 0.689 (gene-protein), 0.632 (protein-miRNA), combined with the between-omics shrinkage factor of 5, resulting in the scaling factor of 0.192 (gene-miRNA), 0.138 (gene-protein), 0.126 (protein-miRNA). After the 5-fold cross-validation, the optimal penalty parameters for molecular profiles are determined to be 0.1 (gene), 0.2 (miRNA), and 0.5 (protein), yielding a total test canonical correlation of 0.799 (normalized scaling factors such that they sum up to 1), and the scaled prediction error of 0.521.

Following this, the complete SmCCNet algorithm is applied with the identified parameters. A subsampling scheme is utilized, selecting 70% of features per iteration for genes and miRNAs and 90% for proteins with 100 iterations to construct the global similarity matrix. Hierarchical clustering with a cut height of 0.995 and a network pruning algorithm set to retain networks between 10 and 100 nodes in size are used to extract the final network modules. The robustness and relevance of the networks are summarized using the NetSHy network summarization score.

After executing the SmCCNet algorithm, we identified five final multi-omics subnetworks (Table 1). Among these, network module 3 demonstrated the strongest association with survival time. Visualization of network module 3 (Figure 7a) through the Shiny application revealed a hub structure centered on the protein *PCNA*, which has strong connections to the miRNAs *miR-150. PCNA* has been studied as a potential biomarker for breast cancer [20], while *miR-150* is known to promote breast cancer growth by targeting the pro-apoptotic purinergic receptor [21]. Despite this, the interaction between *PCNA* and *miR-150* in breast cancer has been minimally studied, leading to a potential area for future validation. Additionally, the correlation heatmap (Figure 7b) indicates strong correlations among molecular features in network module 3, which indicates the efficacy of our hierarchical clustering algorithm in grouping highly correlated molecular features that are significantly associated with the phenotype of interest. Notably, the heatmap reveals an almost perfect correlation among *miR-150, miR-142*, and *miR-146a*, which are closely connected to *PCNA*. This connection hints at a possible immune-related pathway involving *miR-150* and *miR-146a*, particularly their time-dependent relationship with T-cell differentiation [21]. Furthermore, the 3D NetSHy PC scatter plot (Figure 7c) demonstrates that patients with lower survival times tend to be located in the upper part of the 3D space, while those with higher survival times are in the lower part, which suggests that network module 3 could potentially have high predictability of survival time in breast cancer patients. The NetSHy loading plots (Figure 8a-c) reveal that network connections oriented around *PCNA* predominantly influence the first principal component (PC), while *TSC1* -oriented connections play a major role in both the second and third PCs. Notably, the third PC (PC3) exhibits the highest correlation with survival time, with a correlation coefficient (*ρ*) of -0.301. Intriguingly, *TSC1* by itself shows a relatively modest correlation with survival time (*ρ* = -0.119). However, its network connections with other molecular features enhance this association threefold. This implies the importance of further investigating the interactions between *TSC1* and other molecular features within the network, such as *miR-142*, to better understand breast cancer’s biological mechanism.

**Table 1.**
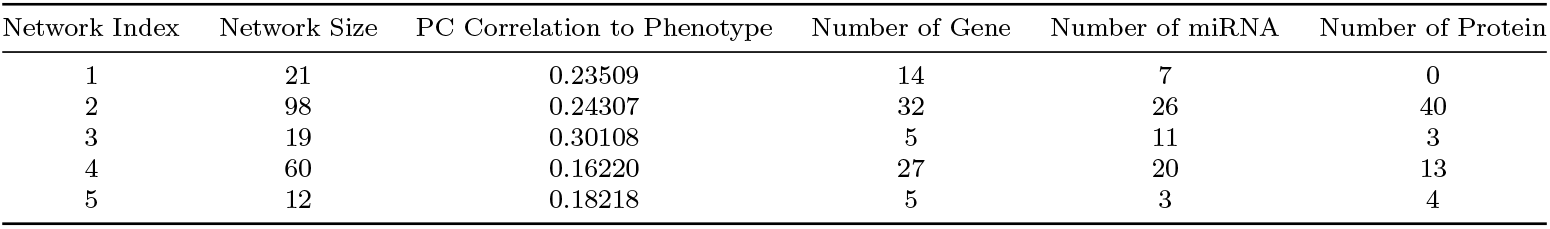
Summary of final subnetwork information for survival time, with information of network index, network size, highest NetSHy score correlation to survival time, number of genes, number of miRNAs, and number of proteins.

**Fig. 7.**
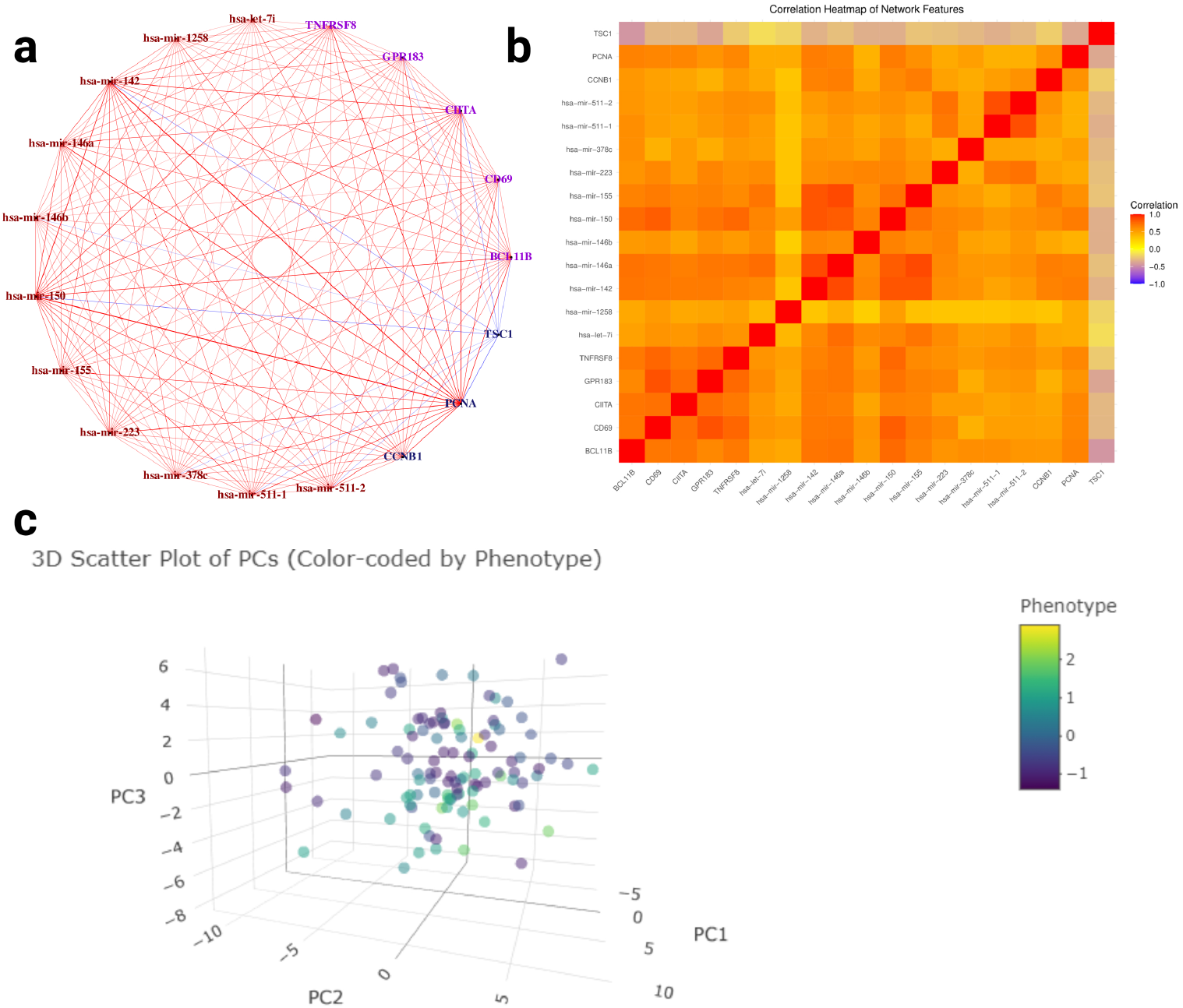
Multi-omics SmCCNet Result for Survival Time. Multi-omics SmCCNet subnetwork result for TCGA breast cancer data with respect to patient’s survival time (subnetwork 3). **a**: Multiomics network with respect to survival time. Purple nodes are genes, brown nodes are miRNAs, and dark blue nodes are proteins. Red edges represent positive association between two nodes, and negative edges represent negative association between two nodes. The color depth and edge width represent the strength of association between two nodes (edges are filtered based on a Pearson’s correlation threshold of 0.3); **b**: the correlation heatmap between all subnetwork molecular features; **c**: 3D scatter plots of subjects based on the first 3 NetSHy PC, with each subject color-coded by survival time (survival time are centered and scaled).

**Fig. 8.**
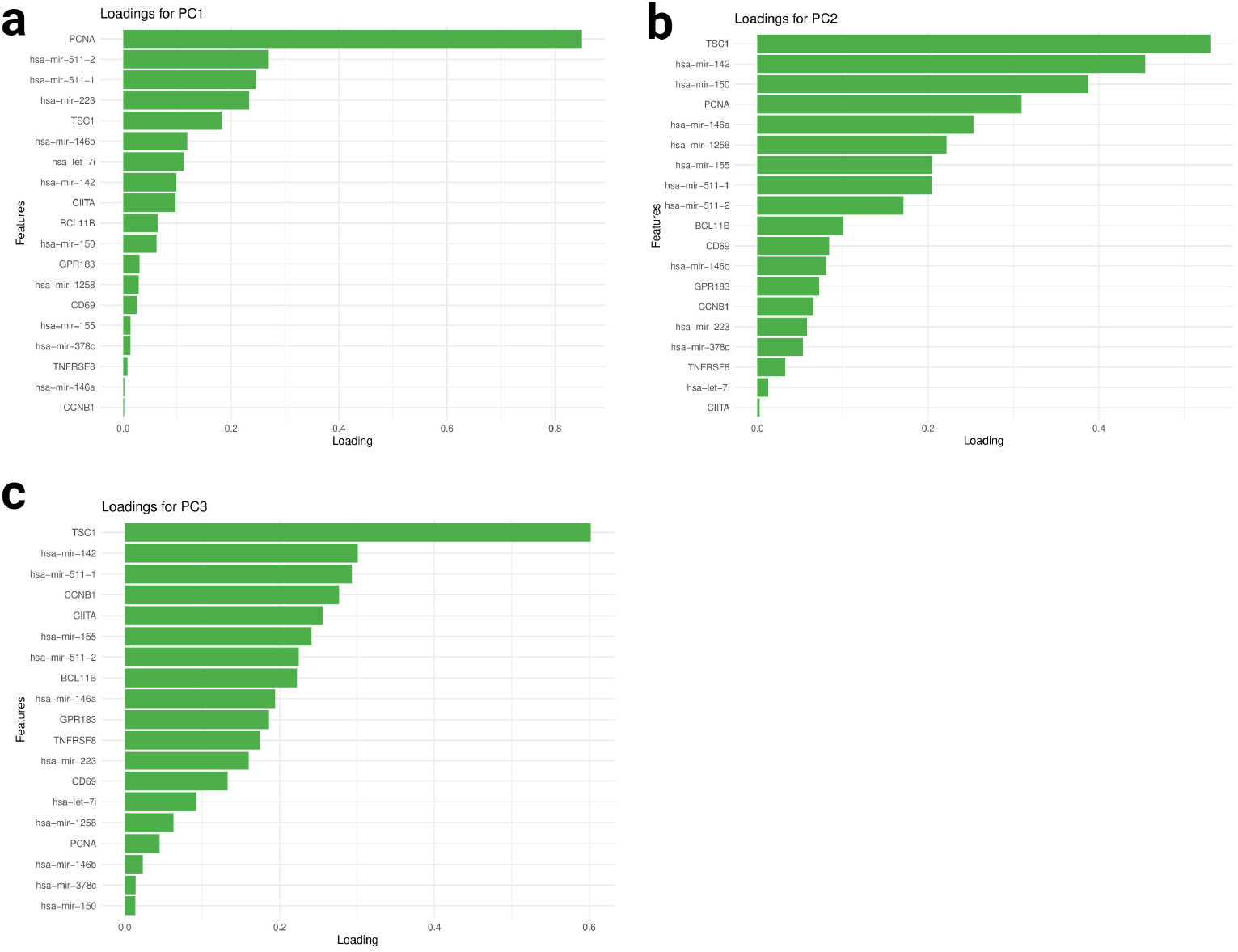
Final Subnetwork NetSHy loadings with respect to survival time. The NetSHy sumamrization loadings of all the final subnetwork features based on subnetwork 3. with panel **a, b**, and **c** represent PC1, PC2, and PC3 respectively

### 3.2 Multi-omics with Binary Phenotype

In the TCGA breast cancer example with a binary phenotype (survival status), the analysis can be run with the following code (assuming all X (-omics data list) and Y (survival status) are preprocessed):

~~~
result ← fastAutoSmCCNet (X = X, Y = **as.factor** (Y),
                          K = 5, subSampNum = 100,
                          DataType = **c** ( ‘Gene’, ‘miRNA’, ‘RPPA’ ),
                          CutHeight = 0.995,
                          summarization = ‘NetSHy’,
                          BetweenShrinkage = 5,
                          seed = 123456,
                          EvalMethod = ‘auc’,
~~~

In the first phase of the SmCCNet algorithm, 5-fold cross-validation is used to optimize the sparsity penalty for each molecular profile and determine the best scaling factors. We consider the SmCCA penalty parameter for each molecular profile ranging from 0.5 to 0.9, with increments of 0.1, and the SPLSDA penalty parameter ranging from 0.5 to 0.9, with increments of 0.1 as well, resulting in 625 combinations. AUC score is used to identify the optimal penalty parameters. Same as before, the preliminary CCA canonical correlation between -omics data is 0.960 (gene-miRNA), 0.689 (gene-protein), 0.632 (protein-miRNA), combined with the between-omics shrinkage factor of 5, resulting in the scaling factor of 0.192 (gene-miRNA), 0.138 (gene-protein), 0.126 (protein-miRNA) for the SmCCA step (exclude phenotype). The relative importance of the between-omics relationship and the omics-phenotype relationship is set to 5, meaning that the omics-phenotype relationship in the model is 5 times as important as the between-omics relationship in the network construction step. After the 5-fold cross-validation, the optimal penalty parameters for SmCCA are determined to be 0.5 (gene), 0.7 (miRNA), and 0.5 (protein); and the optimal penalty parameter for SPLSDA is 0.9, yielding validation AUC score of 0.709. Since SmCCNet is an algorithm that has the balance between molecule interaction and phenotype prediction, and survival status is an outcome that is less associated with -omics profiles, 0.709 is considered a good validation AUC score.

Similarly, as in the quantitative phenotype example, the complete SmCCNet algorithm is applied with the optimal parameters. A subsampling scheme is utilized, selecting 70% of features per iteration for 100 iterations to construct the global similarity matrix. Hierarchical clustering with a cut height of 0.995 and a network pruning algorithm set to retain networks between 10 and 100 nodes in size are used to extract the final network modules. The robustness and relevance of the networks are summarized using the NetSHy network summarization score.

After executing the SmCCNet algorithm, we identified four final multi-omics subnetworks (Table 3). Among these, network module 3 exhibited the strongest association with survival time. Visualization of network module 3 (Figure 9a) through our Shiny application shows that there is less network connectivity after edge filtering based on Pearson’s correlation (threshold = 0.3). After edge filtering, *SLC40A1* has a relatively higher network connectivity. Malignant breast cancer cells modulate their iron metabolism by downregulating the iron exporter gene*SLC40A1* to accommodate their high demand for iron [22]. Additionally, the correlation heatmap (Figure 9b) has a weaker signal than the survival time network, but still demonstrates some high correlation between network molecular features. For instance, there is a strong negative correlation between *TLX1NB* and *SIDT1*. While there is no established study confirming the biological association between *TLX1NB* and *SIDT1* in the context of breast cancer, future studies can be conducted to validate their association. Furthermore, the 3D NetSHy PC scatter plot (Figure 9c) demonstrates that patients with survival status tend to be located in the left part of the 3D space, while those with death status are in the right part, which suggests that network module 3 has strong capability in classifying survival and death case in breast cancer patients, which also implies this network is highly associated with breast cancer.

**Table 2.**
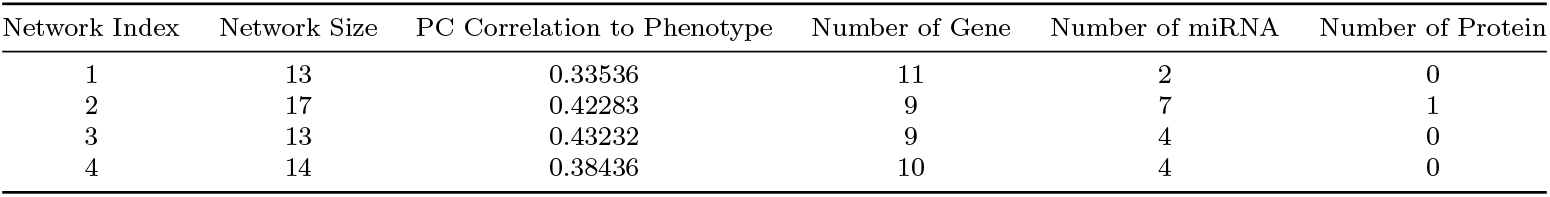
Summary of final subnetwork information for survival time, with information of network index, network size, highest NetSHy score correlation to survival time, number of genes, number of miRNAs, and number of proteins.

**Table 3.**
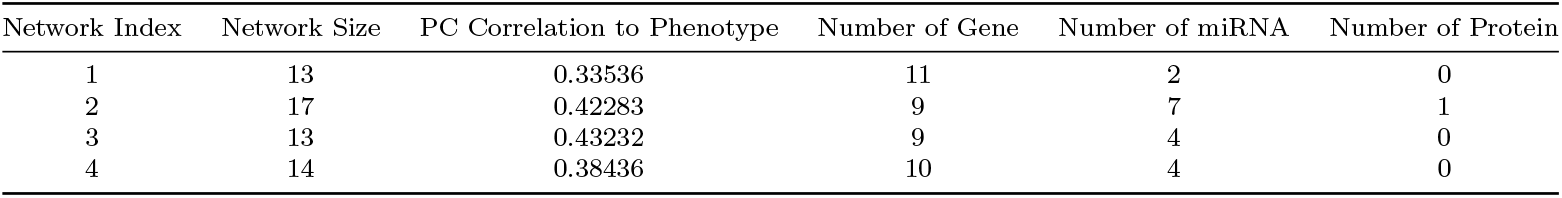
Summary of final subnetwork information for survival time, with information of network index, network size, highest NetSHy score correlation to survival time, number of genes, number of miRNAs, and number of proteins.

**Fig. 9.**
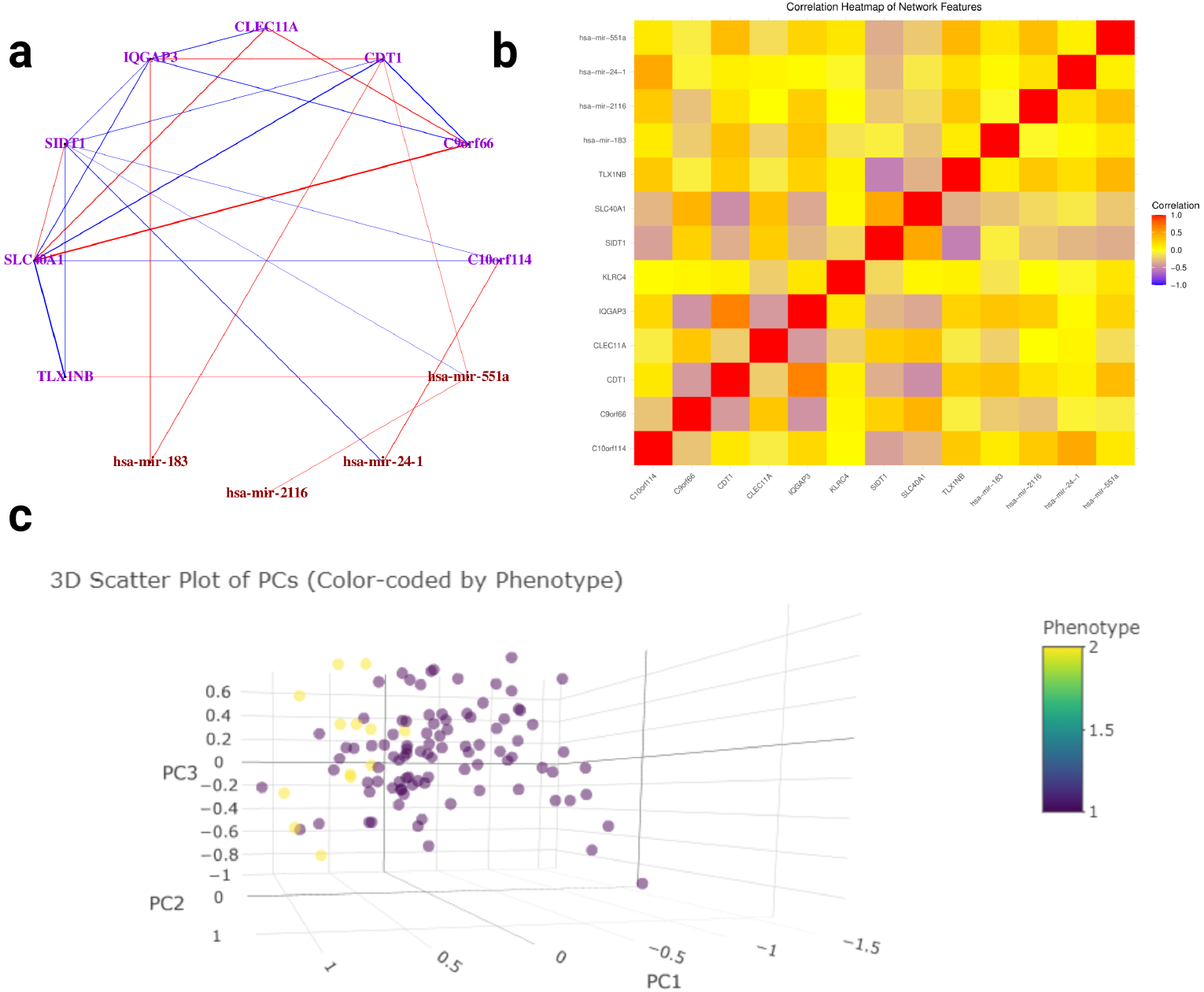
Multi-omics SmCCNet Result for Survival Status. Multi-omics SmCCNet subnetwork results for TCGA breast cancer data with respect to patient’s survival status (subnetwork 3). **a**: Multi-omics network with respect to survival time. Purple nodes are genes, brown nodes are miRNAs, and dark blue nodes are proteins. Red edges represent positive association between two nodes, and negative edges represent negative association between two nodes. The color depth and edge width represent the strength of association between two nodes (edges are filtered based on a Pearson’s correlation threshold of 0.3, after edge filtering, not all network nodes are presented in the network); **b**: the correlation heatmap between all subnetwork molecular features; **c**: 3D scatter plots of subjects based on the first 3 NetSHy PC, with each subject color-coded by survival status.

The NetSHy loading plots (Figure 10a-c) reveal that network connections oriented around *KLRC4* predominantly influence the first and the second principal component (PC), while *miR-24-1* -oriented connections play a major role in both the third PCs. Interestingly, *KLRC4* is filtered out after the stringent Pearson’s correlation edge filtering as shown in Figure 9a, suggesting that it does not have strong interaction with other network molecular features, but it is still influencing the network in some other ways. Additionally, the first PC (PC1) exhibits the highest correlation with survival status, with a biserial correlation coefficient (*ρ*) of -0.432. Interestingly, The highest feature-phenotype correlation is only 0.299, which suggests that adding network interaction improves the prediction on patients’ survival status compared to individual molecular features.

**Fig. 10.**
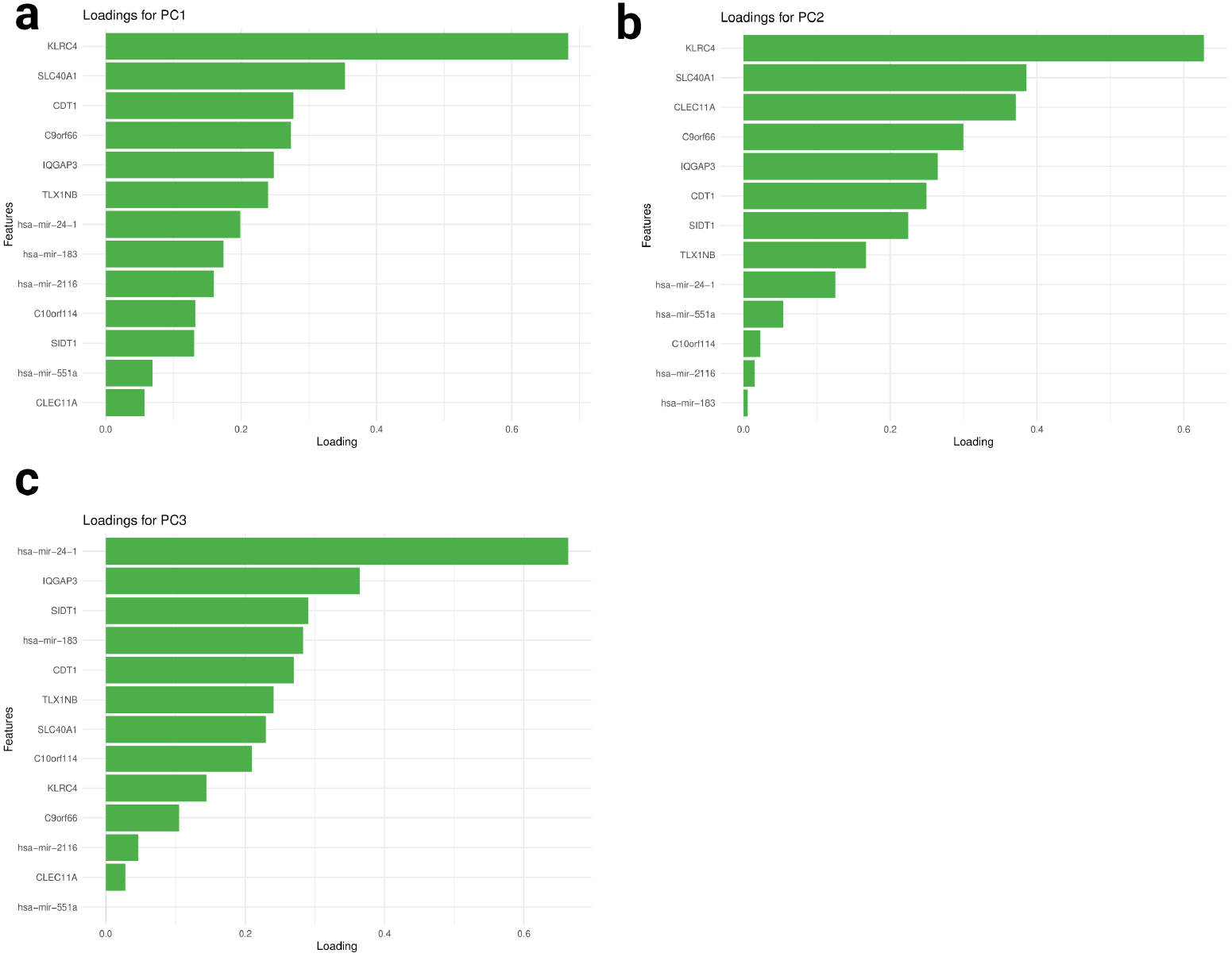
Final Subnetwork NetSHy loadings with respect to survival status. The NetSHy sumamrization loadings of all the final subnetwork features based on subnetwork 3. with panel **a, b**, and **c** represent PC1, PC2, and PC3 respectively.

## 4 Conclusion

The second-generation SmCCNet is a powerful and comprehensive tool for multi-omics network inference with respect to a quantitative or binary variable (e.g., an exposure or phenotype for a complex disease). This upgraded tool incorporates numerous new features including generalization to single or multi-omics data, a novel algorithm for single/multi-omics data with binary phenotype, an automated pipeline to streamline the algorithm with a single line of code, a network pruning algorithm, a topology-based network summarization method, a new network visualization tool, and much more. Additionally, compared to the first-generation SmCCNet, this new version sub-stantially reduces the computational time by up to 1000 times, and the end-to-end pipeline can be set up easily with either a manual form for more specific parameter control, or through the new automated version. In the future, more features such as time-to-event data and longitudinal data will incorporated into the pipeline.

## 5 Acknowledgements

We express our sincere gratitude to Wen (Jenny) Shi and Laura M Saba for their contributions to the development of the first-generation SmCCNet.

## 8 Data Availability

The TCGA breast cancer data used in example section is available at: http://linkedomics.org/data_download/TCGA-BRCA/.

## Footnotes

1 We drop subsampling from step I, which increases the computational speed by 100 - 1000 times

2 https://cran.r-project.org/web/packages/SmCCNet/vignettes/SmCCNet_Vignette_MultiOmics.pdf

3 Penalty terms in SmCCA is in the opposite direction of the SPLSDA, in SmCCA, a higher value of penalty term implies a less stringent sparsity penalty.

4 Example of a more flexible multi-omics SmCCNet pipeline can be found in package vignette https://cran.r-project.org/web/packages/SmCCNet/vignettes/SmCCNet_Vignette_MultiOmics.pdf

